# Optogenetic determination of dynamic and cell-type-specific chloride equilibrium potentials

**DOI:** 10.1101/2023.05.08.539831

**Authors:** Richard J. Burman, Tara Diviney, Alexandru Călin, Gemma Gothard, Jean-Sébastien M. Jouhanneau, James F.A. Poulet, Arjune Sen, Colin J. Akerman

## Abstract

Optogenetics has revolutionized neurobiological research by providing tools for modulating neuronal activity. As these tools utilise light-activated ion fluxes, they afford new opportunities to examine the nature of transmembrane ion gradients. Traditional investigation into the equilibrium potential for chloride (E_Cl_) has been limited to studying endogenous chloride-permeable receptors. Here we demonstrate the utility of using a light-activated chloride channel, stGtACR2, to probe somatic E_Cl_ in rodent. This agonist-independent optogenetic strategy is validated *in vitro* and *in vivo*, captures differences in E_Cl_ dynamics following manipulations of endogenous chloride fluxes, and reveals distinct resting E_Cl_ across genetically-defined neuronal subpopulations. Using this approach to challenge chloride homeostasis, we uncover cell-specific E_Cl_ dynamics that are supported by the differential expression of endogenous handling mechanisms. Our findings establish an optical method for investigating transmembrane chloride gradients and thereby expand the repertoire of optogenetics.

## Introduction

For any ion species, the relative extracellular and intracellular concentrations establish the transmembrane concentration gradient. An ion’s equilibrium potential (E_ion_) refers to the potential difference across the cell membrane that exactly balances this concentration gradient^1, 2^. Knowledge of the equilibrium potential is fundamental to understanding cellular physiology, as increases in membrane permeability to the ion will cause the membrane potential (V_m_) to move towards E_ion_, with the driving force on the ion reflecting how far V_m_ is from E_ion_. If E_ion_ is below the cell’s V_m_, an ion flux across the membrane would result in hyperpolarization. Whilst if E_ion_ is above the cell’s V_m_, the same increase in membrane permeability will result in depolarization. As well as determining the nature of transmembrane ion fluxes, E_ion_ is a parameter that can vary, exhibiting differences between cell types, changes over extended timeframes such as across development, or short-term fluctuations depending upon recent influxes and effluxes.

For example, as the most abundant anion in the body, the equilibrium potential for chloride (E_Cl_) is determined by the transmembrane concentration gradient for chloride (Cl^-^). Intracellular and extracellular Cl^-^ concentrations have been shown to differ between cells and over a range of timescales, consistent with the idea that E_Cl_ is a parameter that varies^3–8^. In the brain, E_Cl_ is particularly important because fast synaptic inhibition is mediated by the GABA-A receptor (GABA_A_R), which is primarily permeable to Cl^-^^9, 10^. The inhibitory effects of GABA_A_Rs therefore depend on both the membrane potential of the neuron and the E_Cl_, which is the product of a dynamic equilibrium between Cl^-^ extrusion and intrusion processes. Such processes include effluxes mediated by the potassium-chloride co-transporter, KCC2^11, 12^, and influxes such as those mediated by GABA_A_Rs themselves^13, 14^. Previous work has established a rich, interconnected set of cellular systems that dynamically regulate E_Cl_ in both space and time, with fundamental implications for our understanding of intercellular signalling in the brain^10, 15, 16^.

Estimates of E_Cl_ have traditionally been made by activating endogenous Cl^-^-permeable receptors on the cell surface and measuring current flow at different V_m_ values. Typically, this involves exogenous application of an agonist (e.g. focal delivery or uncaging of GABA)^17–20^, or stimulation techniques that elicit the release of a relevant endogenous agonist (e.g. electrical or optical activation of presynaptic GABA-releasing axons)^21–25^. There are, however, a number of limitations associated with inferring E_Cl_ from endogenous Cl^-^-permeable receptors. Firstly, it is difficult to be sure which receptor populations are being activated, which is problematic as E_Cl_ is thought to vary between subcellular compartments^19, 26–29^. It can be challenging to deliver the agonist to the cell of interest in intact preparations and it is often necessary to use pharmacology to isolate the receptor’s response. Also, it is difficult to determine the receptor’s own contribution to E ^30, 31^. Furthermore, there are additional processes that influence attempts to measure dynamic E_Cl_ when relying upon endogenous Cl^-^-permeable receptors. Agonist application can saturate receptors, desensitization processes can alter receptor availability^30, 32, 33^, and, when stimulating axonal release, presynaptic mechanisms can alter how the agonist is delivered to the receptor^34, 35^. For these reasons, there is a need to develop additional methods for determining E_Cl_.

The development and application of optogenetics within modern neuroscience has provided an invaluable and ever-expanding toolkit for modulating neuronal activity. Yet, because these tools utilise light-activated ion fluxes, they also afford an untapped opportunity for examining transmembrane ion gradients. Whilst light-activated Cl^-^ pumps, such as halorhodopsin, have been shown to alter E_Cl_ ^36–38^, a more appropriate strategy for measuring E_Cl_ would be to use a light-activated channel that passively reports the transmembrane Cl^-^ gradient, such as an anion channelrhodopsin (ACR)^39–43^. Within this class of opsins, Guillardia theta anion-conducting channelrhodopsin 2 (GtACR2) has excellent performance characteristics due to its high level of ion selectivity and large photocurrents^41, 44, 45^. Furthermore, Mahn and colleagues^46^ have modified the channel with a motif that restricts expression to the somatic compartment (soma-targeted, stGtACR2).

Here we establish stGtACR2’s use as an agonist-independent strategy for probing E_Cl_ in the mouse brain. We demonstrate that this method can quantify the contribution of endogenous Cl^-^ fluxes to E_Cl_ and can be leveraged to distinguish E_Cl_ in different neuronal subpopulations. By using this approach to challenge Cl^-^ homeostasis, we uncover cell-specific E_Cl_ dynamics that are supported by the differential expression of endogenous handling mechanisms. This optogenetic application generates new opportunities for investigating E_Cl_ using light, in a manner that is compatible with the targeting of defined cell populations and subcellular compartments.

## Results

### Using light-activated stGtACR2 to determine Cl^-^ equilibrium potentials

The light-activated Cl^-^ channel, stGtACR2, can be used to elicit Cl^-^ photocurrents at the cell soma^46^. This enables development of an optogenetic strategy for determining the somatic equilibrium potential for Cl^-^ (E_Cl_) (**Figure 1A**). To establish the feasibility of determining E_Cl_ optically, we expressed stGtACR2 in L2/3 pyramidal neurons of mouse cortex. Adeno-associated virus (AAV) encoding Cre-dependent stGtACR2 fused to the red fluorescent protein, FusionRed, was injected into primary somatosensory cortex (S1) of CaMKIIα-Cre mice, at four weeks of age (**Figure 1B**). Having confirmed that this resulted in the expression of stGtACR2-FusionRed in Cux2-expressing L2/3 pyramidal neurons (**Figure 1C**), acute brain slices were prepared two weeks after AAV injections and stGtACR2-expressing L2/3 pyramidal neurons were targeted for whole-cell voltage-clamp patch-clamp recordings (**Figure 1D**). To evoke stGtACR2 photocurrents, brief pulses of blue light (470 nm laser) were targeted at the recorded neuron. Meanwhile, to generate an independent estimate of E_Cl_ in the same neuron, we used a traditional approach of delivering focal puffs of GABA to activate GABA_A_Rs, which are primarily permeable to Cl^-^^10, 47^. The agonist-evoked GABA_A_R currents were pharmacologically isolated by blocking GABA_B_Rs with CGP55845 (10 μM).

**Figure 1:**
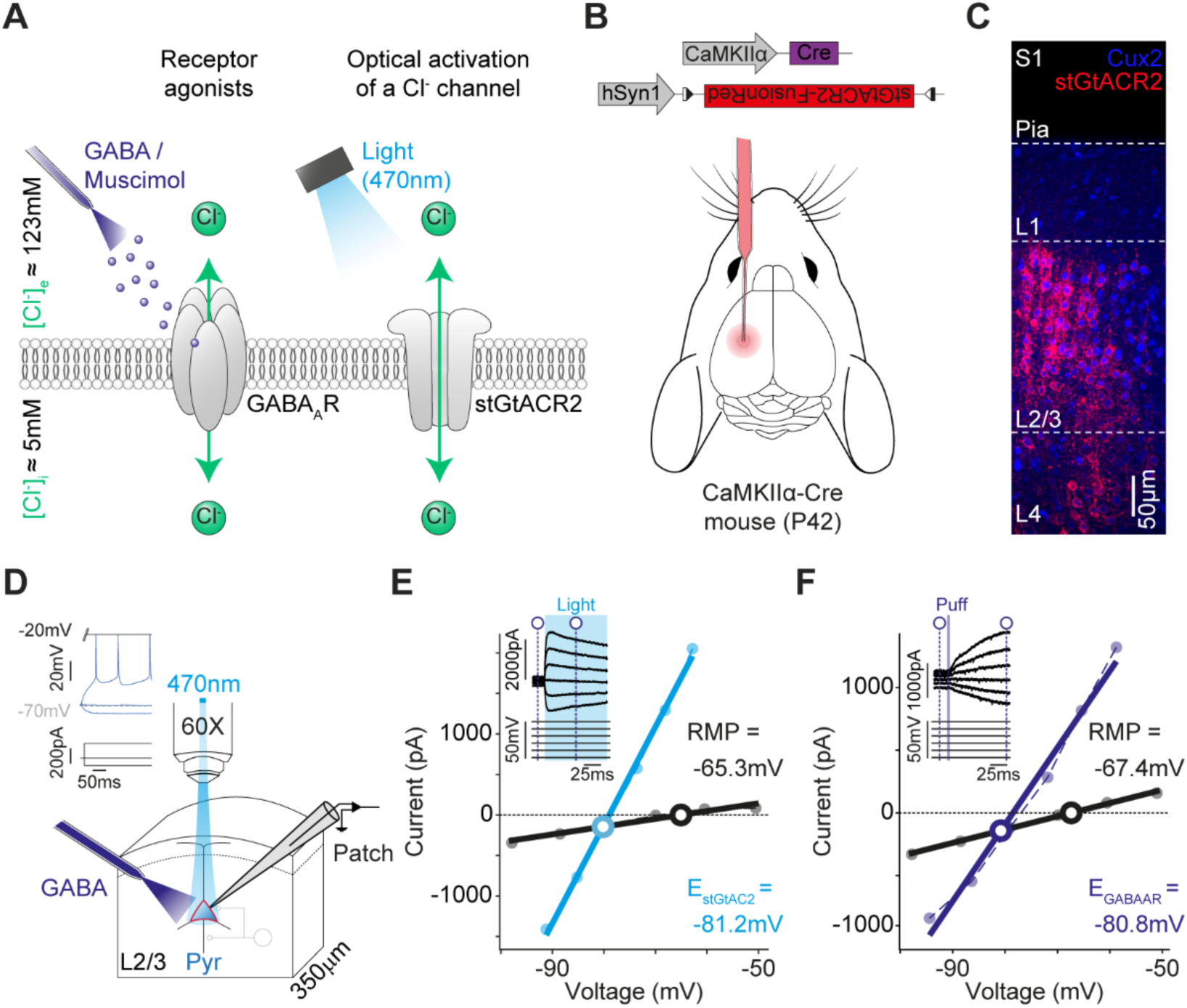
Using light-activated stGtACR2 to determine Cl^-^ equilibrium potentials. (**A**) Schematic showing transmembrane Cl^-^ fluxes elicited by GABA_A_R agonist activation. An optogenetic strategy elicits transmembrane Cl^-^ fluxes via light-activation of the Cl^-^-permeable stGtACR2. (**B**) Cre-dependent stGtACR2-FusionRed was delivered by intracerebral AAV injection into primary somatosensory cortex (S1) of CaMKIIα-Cre mice. (**C**) Confocal image showing stGtACR2-FusionRed expression in Cux2^+^ L2/3 pyramidal neurons at 6 weeks of age, two weeks after injection. (**D**) Experimental setup in which whole-cell patch-clamp recordings were performed from stGtACR2-expressing L2/3 pyramidal neurons. stGtACR2 was activated with blue light pulses (470 nm laser), delivered via the microscope objective. Focal applications of exogenous GABA from a separate pipette were used to activate GABA_A_Rs. Inset, current-clamp recording from an example stGtACR2-expresssing regular spiking pyramidal neuron. (**E**) IV plot from a voltage-clamp step protocol shows the stGtACR2-induced response (cyan, 100 ms light) and baseline current (black) at different holding potentials. The equilibrium potential for both stGtACR2 (E_stGtACR2_) and the baseline current (equivalent to the resting membrane potential, RMP) are indicated. Inset shows raw traces and indicates where baseline and stGtACR2 currents were measured (dashed vertical blue lines). (**F**) IV plot from the same neuron as in ‘E’ showing the GABA-evoked response (purple) and baseline current (black) at different holding potentials, from which E_GABAAR_ and RMP were determined.

The equilibrium potential for stGtACR2 (E_stGtACR2_) was calculated by measuring the amplitude of the light-activated stGtACR2 photocurrent elicited whilst clamping the neuron at different holding potentials (**Figure 1E**). This enabled us to generate separate current-voltage (IV) plots for the baseline holding current and for the total holding current recorded during stGtACR2 activation, whose point of intersection represents E_stGtACR2_ (**Figure 1E**). Meanwhile, an independent estimate of E_Cl_ was made by measuring the amplitude of agonist-evoked GABA_A_R currents elicited whilst clamping at different holding potentials, and inferring the equilibrium potential of the GABA_A_R (E_GABAAR_) in an analogous manner (**Figure 1F**). These recordings revealed similar estimates of E_Cl_ in the same neuron (**Figure 1E-1F**). To explore whether the relationship between E_stGtACR2_ and E_GABAAR_ was maintained across a range of intraneuronal Cl^-^ concentrations, a series of recordings were performed using one of three different intracellular pipette solutions (4 mM, 20 mM or 70 mM Cl^-^), such that the intraneuronal Cl^-^ concentration was altered by dialysis. Across a population of neurons (n = 24 neurons from 7 mice), E_stGtACR2_ and E_GABAAR_ were found to be significantly correlated with one another (p < 0.0001, Pearson correlation coefficient; **Figure 2**), supporting the idea that E_stGtACR2_ can be used to estimate somatic E_Cl_.

**Figure 2:**
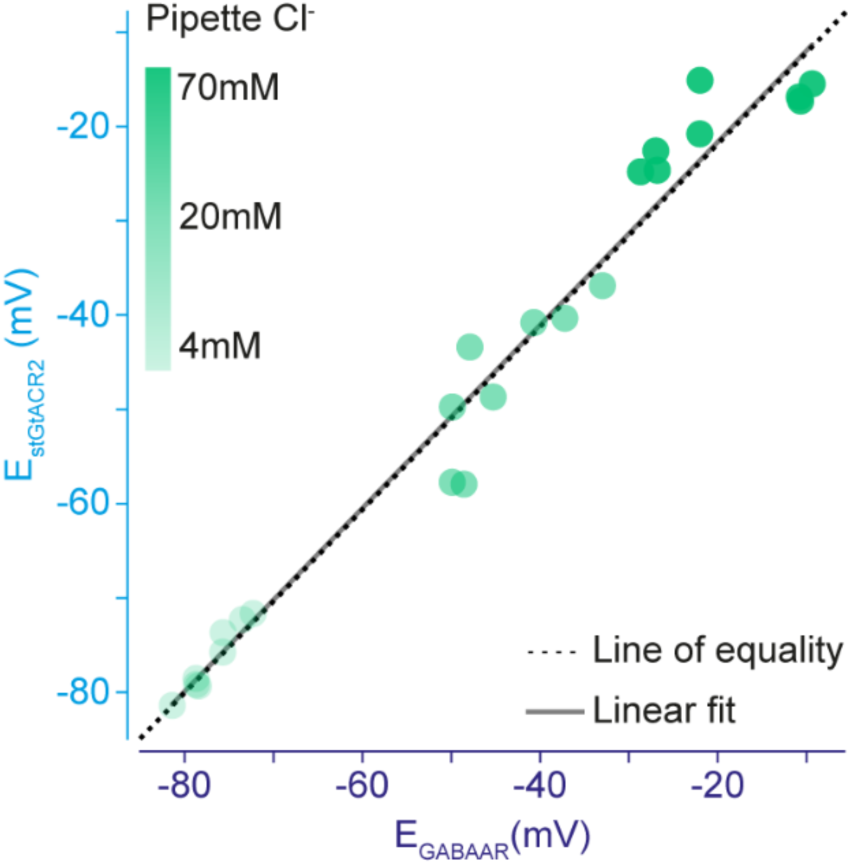
stGtACR2 reports transmembrane Cl^-^ equilibrium potentials. Across a population of neurons using a range of intracellular pipette Cl^-^ concentrations, the equilibrium potential of stGtACR2 (E_stGtACR2_) and the equilibrium potential of agonist-activated GABA_A_Rs (E_GABAAR_) were highly correlated (linear fitted line, black solid line, r = 0.99, n = 24 neurons from 7 mice, p < 0.0001, Pearson correlation coefficient) and approached equality (dashed black line).

### *In vivo* optogenetic measurements of E_Cl_

A significant advantage of determining E_Cl_ optically is that it eliminates the requirement to deliver a receptor agonist, which can be technically challenging, particularly in intact tissue. To illustrate this, we asked whether stGtACR2 could be used to determine E_Cl_ *in vivo* in the intact mouse cortex, by combining two-photon targeted recordings with optical activation of stGtACR2. To optimize the tissue for two-photon imaging, *in utero* electroporation was used to express stGtACR2 in L2/3 pyramidal neurons of S1 (**Figure 3A**). The cytosolic fluorescent reporter protein tdTomato was co-expressed to aid in the screening of the resulting pups (see Methods), and to facilitate identification of the transfected neurons in the mature cortex (**Figure 3B**). Once the animals had reached 6 weeks of age they were anaesthetized with urethane and two-photon targeted *in vivo* voltage-clamp patch-clamp recordings were performed from stGtACR2-expressing L2/3 pyramidal neurons (**Figure 3C**). Alexa 594 dye was added to the patch pipette solution to visualize the pipette under 820 nm light, whilst tdTomato was visualized under 930 nm light (**Figure 3D**).

**Figure 3:**
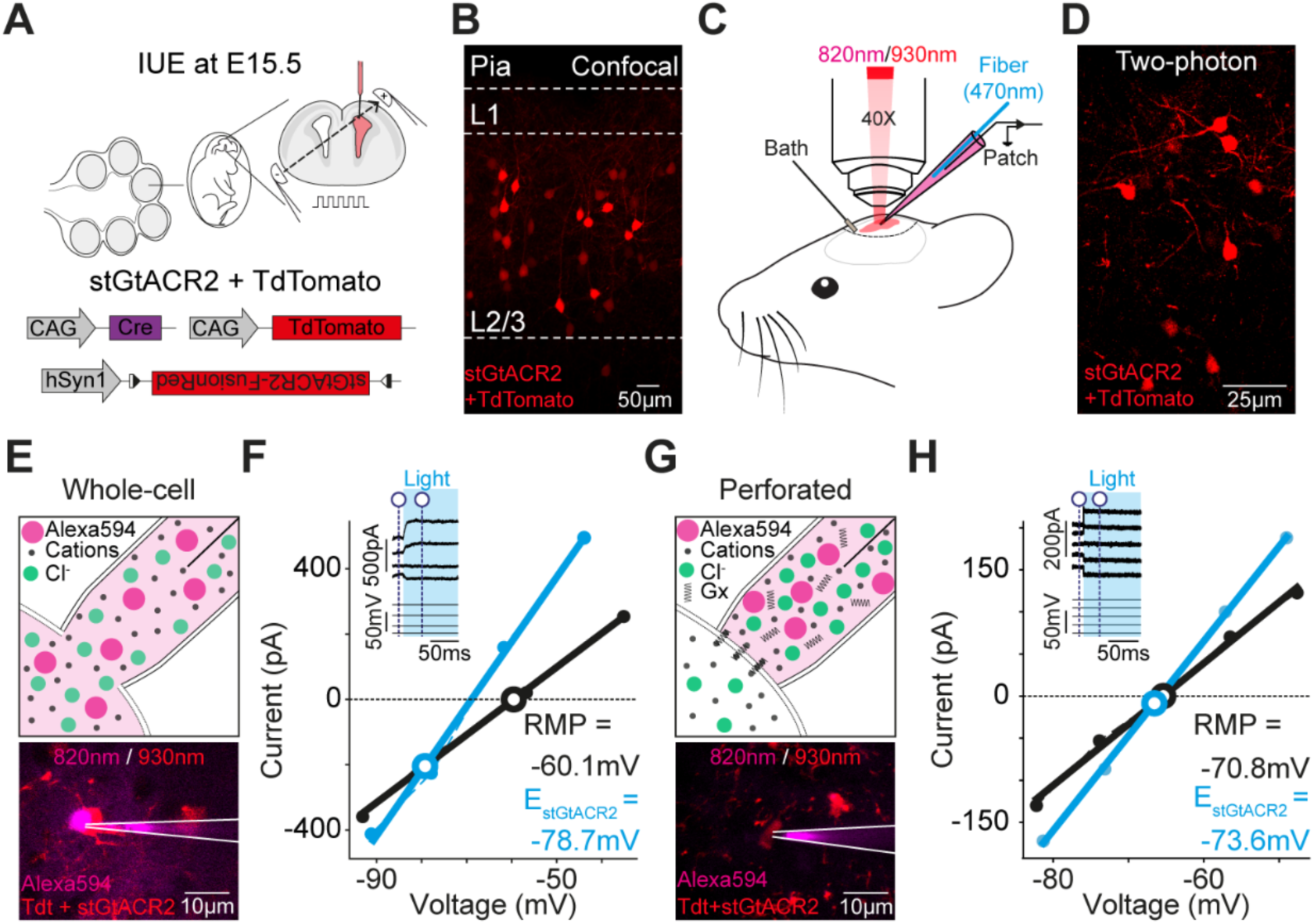
Optogenetic determination of E_Cl_ *in vivo*. (**A**) In utero electroporation (IUE) was used to co-express stGtACR2 and tdTomato in L2/3 pyramidal neurons of mouse S1 cortex. (**B**) Confocal image from S1 at 6 weeks of age. TdTomato facilitated identification of expressing neurons. (**C**) Experimental setup for two-photon targeted patch-clamp recordings. L2/3 pyramidal neurons co-expressing stGtACR2 and tdTomato were visualized at 930 nm. Patch pipettes containing Alexa 594 dye were visualized at 820 nm. Light pulses were delivered via an optic fiber inserted within the patch pipette. (**D**) *In vivo* two-photon image of tdTomato-expressing L2/3 pyramidal neurons. (**E**) Example image of an *in vivo* whole-cell patch-clamp recording (pipette solution, 4 mM Cl^-^). The Alexa 594 dye (magenta) diffuses from the pipette (white lines) into the recorded L2/3 pyramidal neuron. (**F**) IV plot from *in vivo* voltage-step protocol in whole-cell mode shows stGtACR2-induced response (cyan) and baseline current (black) at different holding potentials. Inset shows raw traces and indicates where baseline and stGtACR2 currents were measured (dashed vertical blue lines). The equilibrium potential for stGtACR2 (E_stGtACR2_) and the baseline current (RMP) are indicated. (**G**) Example image of an *in vivo* gramicidin (Gx) perforated patch-clamp recording from a L2/3 pyramidal neuron. Alexa 594 dye does not enter the neuron due to the perforated status of the patch. (**H**) IV plot from *in vivo* gramicidin voltage-step protocol shows stGtACR2-induced response (cyan) and baseline current (black) at different holding potentials.

Initially, recordings performed in whole-cell configuration (**Figure 3E**) confirmed that it was possible to combine light-activation of stGtACR2 currents with voltage-step protocols at different holding potentials, thereby generating estimates of E_stGtACR2_ *in vivo* (**Figure 3F**). Subsequent recordings incorporated the cation-selective perforating agent, gramicidin, into the patch pipette solution, so as to preserve the neuron’s native transmembrane Cl^-^ gradients (**Figure 3G**). We combined our published protocol for performing gramicidin perforated patch-clamp recordings *in vivo*^48, 49^ with two-photon targeting of stGtACR2-expressing neurons. Perforation via gramicidin was confirmed by monitoring the lack of Alexa 594 dye diffusion into the neuron, and from a decrease and then stabilization in series resistance following gigaseal formation (**Figure 3G**). These recordings revealed that it was also possible to estimate resting E_Cl_ under undisturbed conditions *in vivo,* by combining light-activated stGtACR2 currents with voltage-step protocols (**Figure 3H**).

### Optogenetic determination of E_Cl_ dynamics

Transmembrane Cl^-^ fluxes can alter [Cl^-^]_i_, meaning that rather than being static, E_Cl_ is a dynamic parameter that reflects the recent history of the cell. For example, Cl^-^ influxes in neurons cause transient increases in [Cl^-^]_i_ that lead to depolarizing shifts in E_Cl_^36, 50, 51^. The rate at which E_Cl_ subsequently recovers is reflective of the neuron’s capacity to extrude Cl^-^ ^29, 36, 38, 52, 53^. An optogenetic strategy for quantifying E_Cl_ dynamics has the potential to avoid confounds associated with agonist-based techniques, including controlling the amount of agonist, the activation of different receptor pools, and agonist-dependent changes to receptors (**Figure 4A**). Furthermore, by capitalizing on the temporal control afforded by light-activated channels, there is the potential to quantify a cell’s E_Cl_ dynamics and Cl^-^ extrusion processes in real-time, rather than having to infer across separate agonist applications. To investigate this possibility, we performed whole-cell patch-clamp recordings from stGtACR2-expressing L2/3 pyramidal neurons in acute brain slices (**Figure 4B**). We first asked whether stGtACR2 activation can induce an intracellular Cl^-^ load that causes a temporary depolarizing shift in E_Cl_. Adopting a protocol similar to that used by Raimondo and colleagues^36^, we activated stGtACR2 continuously for 1 s whilst depolarizing the membrane potential to increase the driving force for Cl^-^ to enter the neuron (**Figure 4C**). This resulted in a robust and transient Cl^-^ load, as brief stGtACR2 photocurrents switched in polarity, such that outward currents before the Cl^-^ load, became inward immediately after the optical Cl^-^ load, but then decreased in amplitude, and reversed to being outward again over a period of tens of seconds (**Figures 4C** and **4D**).

**Figure 4:**
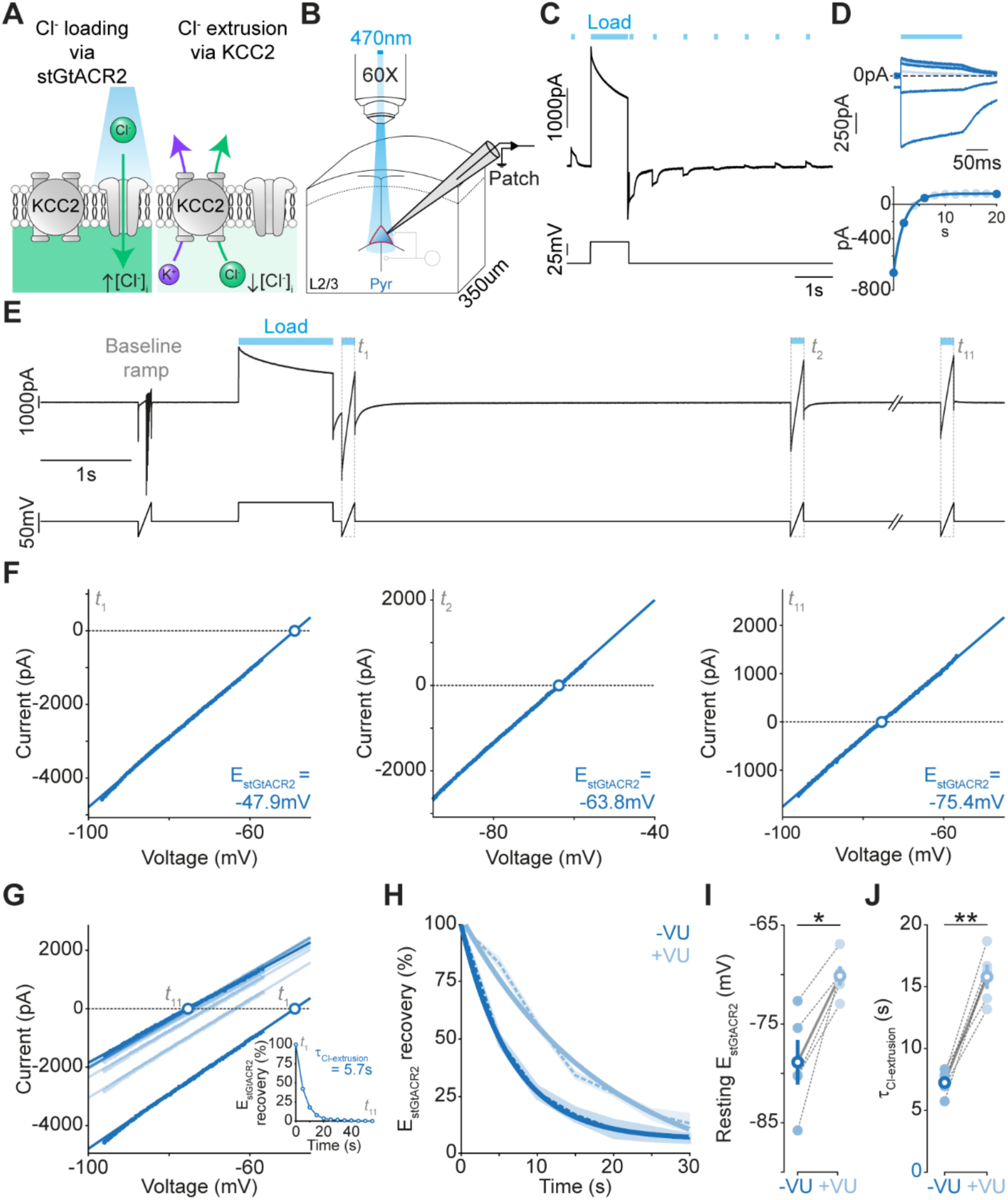
Optogenetic determination of E_Cl_ dynamics. (**A**) Schematic showing intracellular Cl^-^ being raised (‘loaded’) during stGtACR2 activation, and then extruded by KCC2. (**B**) Whole-cell patch-clamp recordings (pipette solution, 4mM Cl^-^) were performed from stGtACR2-expressing L2/3 pyramidal neurons in acute brain slices from S1. (**C**) A neuron experienced a Cl^-^ load by pairing long stGtACR2 activation (1 s light pulse, ‘Load’) with membrane potential depolarization (to -20 mV), to increase the driving force for Cl^-^ to enter the neuron. Following the Cl^-^ load, delivering short light pulses (100 ms) every second revealed that the stGtACR2-current had flipped from being an outward current, to transiently becoming an inward current, which then gradually returned to being an outward current. (**D**) Data from the recording in ‘C’ showing the overlaid stGtACR2-current responses after the Cl^-^ load (top) and a plot capturing the timescale of recovery of the stGtACR2-current responses (bottom). (**E**) Voltage-clamp protocol used to monitor E_stGtACR2_ dynamics relative to a stGtACR2-mediated Cl^-^ load. The traces represent the recorded current (top) and the estimated holding potential following series resistance correction (bottom). The L2/3 pyramidal neuron experienced a Cl^-^ load by pairing long stGtACR2 activation (1 s light pulse, ‘Load’) with membrane potential depolarization (to -20mV). A series of voltage ramps (each 150 ms duration, labelled t_1_ to t_11_) were then paired with a brief stGtACR2-activation (150 ms light pulse) to measure E_stGtACR2_ dynamics at 5 s intervals over a period of 60 s. To isolate the stGtACR2 current, a baseline ramp current (without light) was subtracted from each stGtACR2-associated ramp current. (**F**) IV plots from data in ‘E’ showing the isolated stGtACR2 current at t_1_, t_2_ and t_11_. The x-axis intercept represents E_stGtACR2_. (**G**) IV plot of isolated stGtACR2 currents from t_1_ to t_11_. Inset, plot of E_stGtACR2_ recovery following the Cl^-^ load. A single exponential fit provides the time constant of Cl^-^ extrusion (τ_Cl-extrusion_). (**H**) E_stGtACR2_ recovery dynamics in L2/3 pyramidal neurons under control conditions (-VU; dark cyan) and when KCC2 was blocked with the selective antagonist VU0463271 (+VU, 10 μM; light cyan). Dashed line indicates the mean, shaded area indicates standard error of the mean (SEM). (**I**) KCC2 block results in an increase in resting E_stGtACR2_ (-VU: -78.86 ± 2.56 mV vs. +VU: -70.12 ± 0.98 mV, p = 0.01, paired t-test, n = 5 neurons from 3 mice). (**J**) KCC2 block results in a slowing in Cl^-^ extrusion (-VU: 7.29 ± 0.41 s vs. +VU: 15.79 ± 0.98 s, p = 0.001, paired t-test, n = 5 neurons from 3 mice). *, p < 0.05. **, p < 0.01.

Having established that stGtACR2 can induce Cl^-^ loads, we designed an optogenetic protocol to quantify E_Cl_ dynamics (**Figure 4E**). To generate frequent measurements of E_stGtACR2_ before and immediately following a Cl^-^ load, IV plots of stGtACR2 photocurrents were produced by subtracting the current response to a control voltage ramp (‘baseline ramp’) from the response to the same voltage ramp delivered during stGtACR2 activation with blue light (**Figure 4E** and **4F**). These repeated measurements revealed that E_stGtACR2_ had become more depolarized immediately following an optical Cl^-^ load, and recovered to hyperpolarized values over a period of seconds. From these real-time dynamic measurements, the neuron’s Cl^-^ extrusion capacity could be defined as the time constant of E_stGtACR2_ recovery dynamics (τ_Cl-extrusion_, **Figure 4G**), which was 7.29 ± 0.41 s under control conditions (n = 5 neurons from 3 mice). To confirm the contribution of KCC2 to these E_Cl_ dynamics, we added the KCC2-specifc antagonist VU0463271 (‘VU’) to the perfusate and repeated the protocol (**Figure 4H**). In addition to an effect upon resting E_stGtACR2_ values (**Figure 4I**), blocking KCC2 resulted in a slowing in the rate of E_stGtACR2_ recovery dynamics (**Figure 4H**), such that the τ_Cl-extrusion_ more than doubled (**Figure 4J**). Thus, our measurements of E_stGtACR2_ provide an effective readout of E_Cl_ dynamics, whilst avoiding the confounders associated with agonist-based techniques.

### Optogenetic discrimination of neuron-specific differences in E_Cl_

Optimal methods for determining E_Cl_ should be able to distinguish cells that differ in their Cl^-^ handling mechanisms. Several reports have suggested that parvalbumin-expressing (PV) fast-spiking interneurons and pyramidal neurons exhibit differences in their Cl^-^ homeostasis, although there has been no consensus on the relative contribution that cation Cl^-^ co-transporters (CCCs) make in these neuron types^29, 54–58^. We took advantage of the Cre-lox recombination system and our optogenetic approach to investigate potential differences in resting E_Cl_ and E_Cl_ dynamics between PV interneurons and pyramidal neurons. We performed intracortical viral injections of Cre-dependent stGtACR2-FusionRed into S1 of PV-Cre transgenic mice (**Figure 5A**). Using this approach, we were able to restrict stGtACR2 expression to PV interneurons (**Figure 5B**), which could then be targeted for gramicidin perforated patch-clamp recordings, to preserve native transmembrane Cl^-^ gradients (**Figure 5C**). Equivalent recordings were performed from pyramidal neurons following stGtACR2 expression in S1 of CaMKIIα-Cre mice (as in **Figure 1B**).

**Figure 5:**
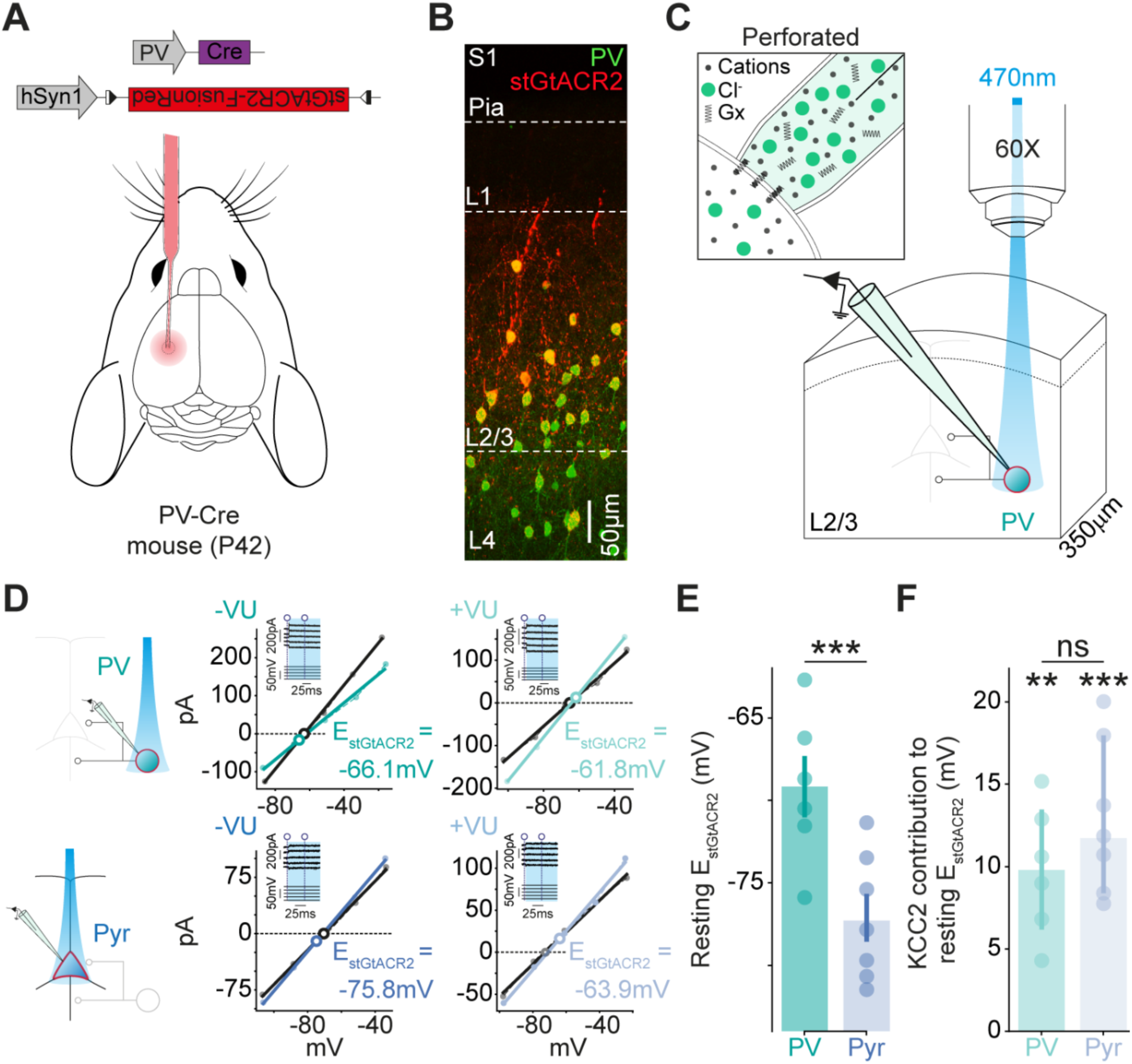
Optogenetic discrimination of neuron-specific differences in E_Cl_. (**A**) Mice expressing Cre recombinase in parvalbumin (PV) interneurons received S1 injections of Cre-dependent stGtACR2-FusionRed AAV. (**B**) Confocal image showing selective expression of stGtACR2-FusionRed (red) in L2/3 immunohistochemically confirmed PV interneurons (green). (**C**) Setup in which gramicidin (Gx) perforated patch-clamp recordings were performed from stGtACR2-expressing PV interneurons. (**D**) Resting E_stGtACR2_ was measured in PV interneurons (turquoise, top) and pyramidal neurons (blue, bottom) using a voltage-clamp step protocol. The resulting IV plots indicate resting E_stGtACR2_ under control conditions (-VU; darker, middle) and when KCC2 was blocked with VU0463271 (+VU; lighter, right). (**E**) Resting E_stGtACR2_ was more depolarized in PV interneurons compared to pyramidal neurons (PV: -69.16 ± 4.54 mV, n = 6 neurons from 3 mice vs. Pyr: -77.12 ± 3.83 mV, n = 7 neurons from 5 mice, p = 0.006, unpaired t-test). Pyramidal neuron data replotted from Figure 4. (**F**) PV interneurons and pyramidal neurons both exhibited a significant contribution of KCC2 to resting E_stGtACR2_ (PV: 9.79 ± 1.62 mV, n = 6 neurons from 3 mice, p = 0.002, one-sample t-test; Pyr: 12.92 ± 1.76 mV, n = 7 neurons from 5 mice, p = 0.0003, one-sample t-test), and there was no detectable difference in the contribution of KCC2 to resting E_stGtACR2_ in the two neuronal populations (p = 0.23, unpaired t-test). **, p < 0.01; ***, p < 0.001; ns, not significant.

Voltage step protocols were used to compare resting E_stGtACR2_ in L2/3 PV interneurons and pyramidal neurons, before and after pharmacologically blocking KCC2 with VU (**Figure 5D**). In line with previous reports^29, 54–58^, this revealed a more depolarized resting E_stGtACR2_ in PV interneurons than pyramidal neurons (**Figure 5E**), thereby confirming that optogenetic E_Cl_ measurements can distinguish between neuron types. VU application caused a depolarizing shift in the resting E_stGtACR2_ of PV interneurons and pyramidal neurons, revealing that KCC2-mediated extrusion contributes to resting E_Cl_ in both neuron types (**Figure 5D** and **5F**). Similar to previous work^29^, the VU-associated shifts in resting E_stGtACR2_ were comparable for the two neuron types, suggesting that the neuron types could not be distinguished in terms of KCC2’s contribution (**Figure 5F**).

### Optogenetic E_Cl_ dynamics reveal neuron-specific KCC2-mediated extrusion

It has been suggested that KCC2 activity is better revealed when a neuron is subjected to a Cl^-^ load, as this challenges the neuron’s capacity to extrude Cl^-53, 59, 60^. To test if optogenetic E_Cl_ measurements can be used to resolve more subtle cell-type differences in KCC2 activity, we used whole-cell patch-clamp recordings to quantify E_stGtACR2_ dynamics in L2/3 PV interneurons and compared these to equivalent recordings in pyramidal neurons (**Figure 6A**). As we had observed in pyramidal neurons (**Figure 4**), E_stGtACR2_ in PV interneurons exhibited depolarized values immediately following a stGtACR2-mediated Cl^-^ load, and then gradually returned to hyperpolarized values over a period of seconds (**Figure 6B** and **6C**). Each PV interneuron’s Cl^-^ extrusion capacity was defined from the time constant of E_stGtACR2_ recovery dynamics (τ_Cl-extrusion_, **Figure 6C**), which was shown at the population level to slow when KCC2 was blocked with VU (**Figure 6D** and **6E**).

**Figure 6:**
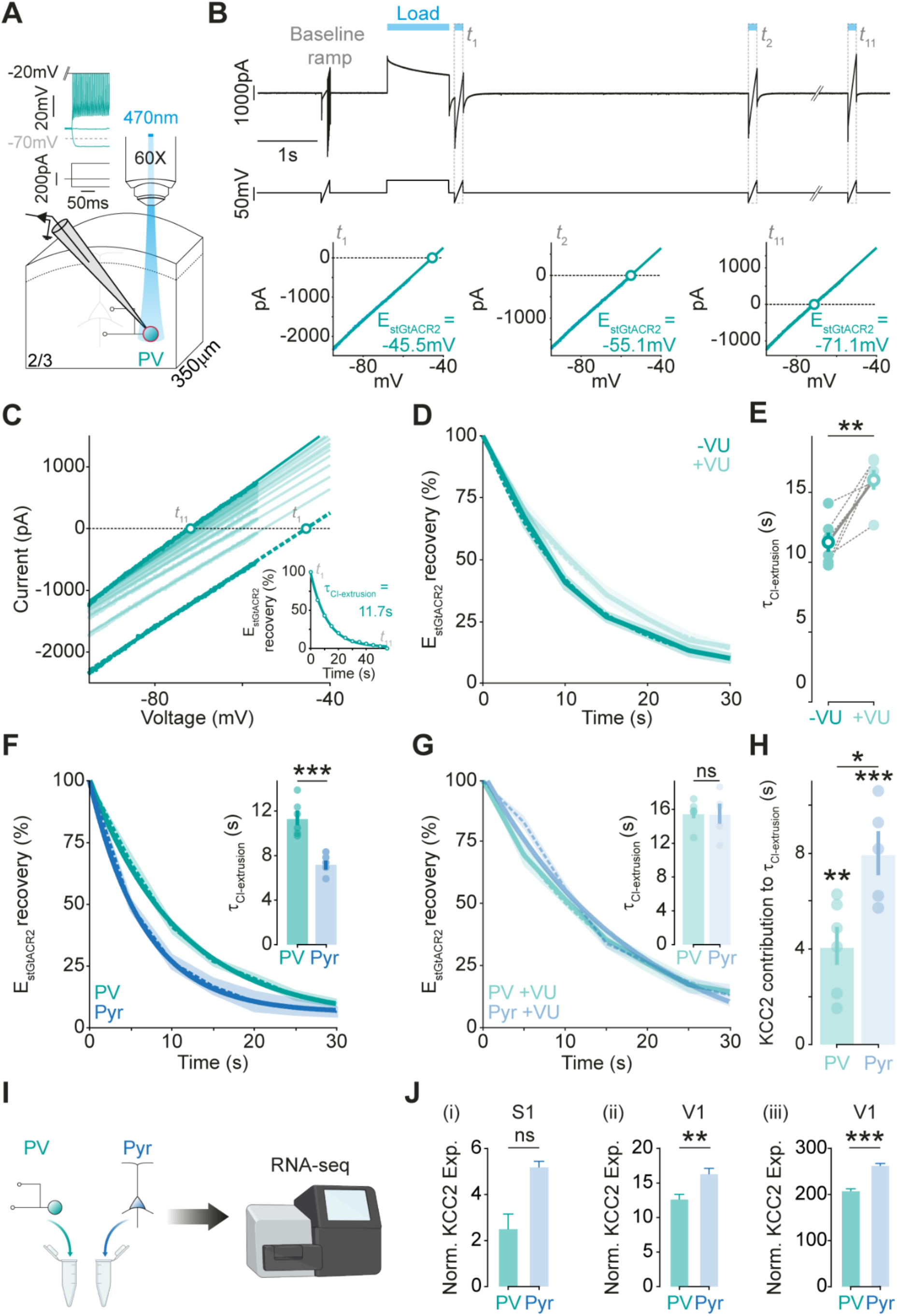
Optogenetic E_Cl_ dynamics reveal neuron-specific differences in KCC2-mediated extrusion. (**A**) Whole-cell patch-clamp recordings (pipette solution, 4 mM Cl^-^) were performed from stGtACR2-expressing L2/3 PV interneurons in S1. Inset, current-clamp recording from an example stGtACR2-expresssing fast-spiking PV interneuron. (**B**) Voltage-clamp ramp protocol used to monitor E_stGtACR2_ recovery following a stGtACR2-mediated Cl^-^ load. Example data from a PV interneuron indicate the recorded current (top), estimated holding potential following series resistance correction (middle), and resulting IV plots (bottom) showing the isolated stGtACR2 current at t_1_, t_2_ and t_11_. Conventions as in Figure 4. (**C**) IV plot of isolated stGtACR2 currents from t_1_ to t_11_ in a PV interneuron. Inset, E_stGtACR2_ recovery following Cl^-^ load. A single exponential fit provides the time constant of Cl^-^ extrusion (′″_Cl-extrusion_). (**D**) E_stGtACR2_ recovery in PV interneurons under control conditions (-VU; dark cyan) and when KCC2 was blocked with the selective antagonist VU0463271 (+VU, 10 μM; light cyan). (**E**) KCC2 block results in a slowing in Cl^-^ extrusion in PV interneurons (-VU: 11.38 ± 0.65 s vs. +VU: 15.57 ± 0.63 s, p = 0.006, paired t-test, n = 6 neurons from 3 mice). (**F**) E_stGtACR2_ recovery following a Cl^-^ load was slower in PV interneurons than pyramidal neurons. Inset, Cl^-^ extrusion time constant (′″_Cl-extrusion_) was greater in PV interneurons (PV: 11.38 ± 0.65 s vs. Pyr: 7.29 ± 0.41 s, p = 0.0007, unpaired t-test). Pyramidal neuron data replotted from Figure 4. (**G**) E_stGtACR2_ recovery and the resulting Cl^-^ extrusion time constant were indistinguishable in PV interneurons and pyramidal neurons after KCC2 block (PV: 15.57 ± 0.63 s vs. Pyr: 15.79 ± 0.98 s, p = 0.84, unpaired t-test). (**H**) Whilst PV interneurons and pyramidal neurons both exhibited a significant contribution of KCC2 to their Cl^-^ extrusion time constant (PV: p = 0.004, one-sample t-test; Pyr: p = 0.001, one-sample t-test), KCC2’s contribution to Cl^-^ extrusion was significantly less in PV interneurons (PV: 4.13 ± 0.79, n = 6 neurons from 3 mice vs. Pyr: 8.01 ± 0.92, n = 5 neurons from 3 mice, p = 0.01, unpaired t-test). (**I**) Meta-analysis of single-cell RNA sequencing data on KCC2 levels in isolated pyramidal neurons and PV interneurons from sensory cortex. (**J**) Normalized KCC2 RNA expression levels are lower in PV interneurons than pyramidal neurons in three separate studies: (i) Zeisel and colleagues^61^ in S1 (PV: 2.54 ± 0.62, n = 10 neurons vs. Pyr: 5.23 ± 0.23, n = 286 neurons, p = 0.07, Kolmogorov-Smirnov test Dunn’s multiple comparison test), (ii) Tasic and colleagues^62^ in primary visual cortex (V1; PV: 9.73 ± 0.59, n = 270 neurons vs. Pyr: 16.40 ± 0.71, n = 756 neurons, p = 0.0073, Kolmogorov-Smirnov test), and (iii) Tasic and colleagues^63^ in V1 (PV: 209.5 ± 3.25, n = 1337 neurons vs. Pyr: 264.3 ± 2.94, n = 7366 neurons, p < 0.0001, Kolmogorov-Smirnov test). *, p < 0.05; **, p < 0.01; ***, p < 0.001.

A direct comparison of these dynamics across cell types revealed clear differences between PV interneurons and pyramidal neurons (**Figure 6F**, **6G** and **6H**). Firstly, the rate of E_stGtACR2_ recovery was slower in PV interneurons than in pyramidal neurons (**Figure 6F**). This cell-type difference was abolished when KCC2 was blocked with VU (**Figure 6G**), and a direct comparison of KCC2’s contribution to τ_Cl-extrusion_ revealed that KCC2 makes a smaller contribution to Cl^-^ extrusion mechanisms in PV interneurons (**Figure 6H**). To corroborate these functional observations, we performed a meta-analysis of single-cell RNA sequencing studies that have distinguished pyramidal neurons and PV interneurons in mouse sensory cortex^61–63^ (**Figure 6I**). After normalizing for total gene expression per cell, we observed that the expression levels of KCC2 were lower in PV interneurons compared to pyramidal neurons (**Figure 6J**). This difference was evident in two independent large-scale datasets from primary visual cortex (V1), and a similar trend was also observed in a dataset from S1 cortex that comprised a smaller number of PV interneurons. Taken together, these results support the conclusion that optogenetic estimates of E_Cl_ dynamics can reveal functionally relevant differences in neuron-specific Cl^-^ extrusion mechanisms.

## Discussion

Here we describe an optogenetic strategy for probing resting and dynamic E_Cl_ in a cell-specific and subcellular-targeted manner. We demonstrate that this approach meets important prerequisites for studying transmembrane Cl^-^ gradients, whilst offering a series of advantages over traditional agonist-based methods. The strategy is validated *in vitro* and *in vivo*, is shown to capture differences in E_Cl_ dynamics following manipulations of endogenous Cl^-^ fluxes, and reveals distinct resting E_Cl_ across genetically-defined neuronal subpopulations. Furthermore, using our optogenetic approach to challenge Cl^-^ homeostasis, we quantify a cell’s E_Cl_ dynamics and uncover cell-specific Cl^-^ extrusion processes that are supported by the differential expression of endogenous handling mechanisms.

Our characterization focused on the use of the soma-targeted, light-activated, Cl^-^ selective channel, stGtACR2^41, 46, 64^. To establish the potential of using stGtACR2 to estimate E_Cl_, E_stGtACR2_ was determined across a range of intracellular Cl^-^ concentrations (4 - 70 mM) and compared to estimates of the equilibrium potential for the GABA_A_R - an endogenous receptor that is primarily permeable to Cl^-^. The strong, positive correlation across a population of neurons supported the conclusion that E_stGtACR2_ reflects E_Cl_, despite potential sources of variation including the location of the two Cl^-^ fluxes in the recorded cells, the modest permeability of the GABA_A_R to bicarbonate^47^, and the extent to which each neuron’s intracellular Cl^-^ had been dialyzed. Further validation came from our measurements of E_stGtACR2_ dynamics and the fact that the time constant for E_stGtACR2_ recovery following a Cl^-^ load (approximately 7.3 s) was close to previous estimates of the time constant for E_Cl_ recovery (approximately 8 s across six separate studies)^11, 36, 50, 51, 53, 65^ - a parameter that has been shown to be invariant to the size of Cl^-^ load^53^. Finally, we also confirmed that E_stGtACR2_ is sensitive to manipulations of Cl^-^ homeostasis in multiple cell types, by pharmacologically blocking KCC2^11,59,66,67.^

One advantage of an optogenetic approach is that it removes technical challenges associated with activating ligand-gated endogenous receptors and therefore lends itself to studies in intact preparations. Accordingly, we show that stGtACR2 can be used to determine E_Cl_ in defined neurons of the intact brain, generating the potential for future experiments that could combine dynamic E_Cl_ measurements and manipulations of Cl^-^-regulatory processes *in vivo*. This application represents an important complement to the use of Cl^-^ imaging approaches, which can provide information on cytosolic Cl^-^ concentration *in vivo*, but do not capture E_Cl_^68–70^. The optogenetic strategy also avoids other limitations associated with inferring E_Cl_ from endogenous Cl^-^-permeable receptors, such as the requirement to use drugs in order to isolate the endogenous Cl^-^ receptor’s response^71, 72^. Such drugs can indirectly alter E_Cl_ by affecting network activity, as well as making it difficult to disentangle the receptor’s own contribution to E_Cl_^30, 31^

Another challenge with agonist-based approaches is that it can be difficult to be sure which receptor populations are being activated. This source of variance is particularly relevant when one considers that E_Cl_ can vary spatially across a neuron and differ between subcellular compartments^19, 26–29^. An attractive feature of stGtACR2 is that it can be targeted to different subcellular compartments, and/or combined with focal light delivery. Whilst our work has characterized a soma-targeted GtACR2, it will therefore be important to explore alternative protein targeting motifs to investigate E_Cl_ in both dendrites and at the axon initial segment. By targeting stGtACR2 to the soma of genetically-defined cell types, we were able to demonstrate that PV interneurons and pyramidal neurons in L2/3 of mouse visual cortex differ in their resting E_stGtACR2_. This supports previous evidence of more depolarized E_Cl_ values in interneurons of the hippocampus and neocortex^29, 55, 73^. More generally, these observations indicate that relatively high baseline somatic E_Cl_ may be integral to interneuron function. For example, a relatively depolarized E_Cl_ in PV interneurons may favour shunting inhibition rather than hyperpolarizing inhibition and enable these interneurons to promote coherent oscillatory network activity^57^. An interesting area for future work will be to use optogenetic approaches to determine E_Cl_ dynamics in the different, genetically-defined, interneuron subpopulations.

In terms of the mechanisms that determine a cell’s transmembrane gradients, we found that KCC2 makes a similar contribution to resting E_Cl_ in both PV interneurons and pyramidal neurons, supporting recent evidence from hippocampus^29^. This would suggest that KCC2 is not responsible for the different resting E_Cl_ between these two neuron types, and that other transmembrane Cl^-^ fluxes contribute. The biophysical prediction is that more subtle differences in a co-transporter’s activity can be revealed under an ionic load, which might better represent a neuron’s state within an active network^16^. In line with this prediction, we capitalized upon the temporal and titratable control of stGtACR2, which make an optical strategy well suited to both challenge and then monitor a cell’s E_Cl_ through rapid and sequential measurements. This enabled us to quantify a cell’s E_Cl_ dynamics and Cl^-^ extrusion processes in real-time, which is difficult with agonist-based approaches^36^. Furthermore, by restricting transmembrane Cl^-^ fluxes to occur only during the light pulse, our optical approach minimizes any alteration to E_Cl_ caused by sustained Cl^-^ fluxes, as observed with receptor agonists^74, 75^. The transient nature of stGtACR2 Cl^-^ fluxes are also less likely to interfere with Cl^-^-sensitive intracellular pathways, such as the With-no-lysine (WNK) kinases that regulate Cl^-^ cotransporter function^76, 77^. Minimizing effects on the cell’s endogenous signaling pathways therefore distinguishes our optogenetic approach from techniques that involve the application of a continuous intracellular Cl^-^ load, via a whole-cell patch pipette for example^59^.

In applying our optogenetic method, we have generated new insights into cell-type-specific somatic E_Cl_ dynamics and their underlying mechanisms. We reveal that PV interneurons exhibit a slower recovery to a transient Cl^-^ load and that this difference in E_Cl_ dynamics is due to a differential contribution by KCC2, as blocking the co-transporter normalized E_Cl_ dynamics across neurons. This was corroborated by single-cell RNA sequencing data from mouse sensory cortex, which reveal higher KCC2 expression in pyramidal neurons than PV interneurons. We therefore propose that our optogenetic strategy generates sensitive estimates of E_Cl_ dynamics, avoiding issues associated with repeated activation of endogenous Cl^-^-permeable receptors.

In conclusion, we establish an optogenetic and agonist-independent strategy for probing E_Cl_. This represents the first example of using a light-activated channel to estimate equilibrium potentials. More generally, this underscores the fact that optogenetic techniques afford opportunities beyond their principal application of modulating neuronal activity. Our findings help to expand the repertoire of optogenetics and encourage the application of other light-activated ion channels in physiological investigations of transmembrane ion gradients and their associated equilibrium potentials.

## Methods

### Intracortical viral injections

To selectively express the soma-targeted light-activated stGtACR2 Cl^-^ channel in excitatory or inhibitory cortical neuronal subpopulations, the Cre-dependent construct pAAV_hSyn1-SIO-stGtACR2-FusionRed packaged in an adeno-associated virus (AAV1, <u>#105677-AAV1</u>, Addgene) was injected into primary somatosensory cortex (S1) of mice homozygous for either CaMKIIα-Cre (Tg(Camk2a-cre)T29-1Stl/J; The Jackson Laboratory) or PV-Cre (B6;129P2-Pvalb^tm1(cre)Arbr^/J; The Jackson Laboratory). Briefly, mice (4 weeks, P28) were anaesthetized with isoflurane (Zoetis) and placed in a stereotaxic frame (Kopf). Analgesic management included intravenous meloxicam (5 mg/kg) and buprenorphine (0.1 mg/kg) as well as intradermal marcain (2 mg/kg) injected into the scalp. Each cornea was protected with ointment (Viscotears) and the animal’s temperature was monitored throughout the procedure. The scalp was shaved (Wahl) and cleaned with a chlorhexidine-based solution (Hibiscrub). A midline incision was made that extended from the interpupillary line to the interaural line and careful dissection was performed to expose the underlying cranium. The craniotomy (diameter less than 1 mm) was located over the left S1 (coordinates in relation to bregma: 3 mm lateral, 1.2 mm posterior) and performed with a dental drill (Foredom). A bevelled glass micropipette (Blaubrand IntraMARK) preloaded with undiluted virus was inserted 300 μm below the surface of the cortex. A total volume of 500 nL was injected at a rate of 33 nL/min. After every 100 nL, the micropipette was raised in 50 μm increments. Following completion of the injection, the micropipette was left in situ for an additional 5 min before it was removed. The scalp was sutured (6-0 Vicryl, Ethicon) and the animal recovered before being returned to its home cage. Following injections, mice were maintained for at least 2 weeks.

### In utero electroporation

Surgery was performed upon timed pregnant female C57BL/6 wild-type mice, with the day of plugging defined as embryonic day 0.5 (E0.5). In utero electroporation (IUE) was performed at E15.5 to maximize targeting of cortical pyramidal neurons in L2/3 ^78^. On the day of the procedure, the pregnant dam was anaesthetized with isoflurane (Zoetis) and given subcutaneous analgesia (5mg/kg meloxicam and 0.1mg/kg buprenorphine). A midline laparotomy was performed and the uterine horns were exposed. DNA plasmids were loaded into a micropipette made from borosilicate glass capillaries (Harvard Apparatus). The DNA plasmids consisted of: Cre recombinase under the Chicken β-actin (CAG) promoter (<u>#13775</u>, Addgene), Cre-dependent stGtACR2-FusionRed under the hSyn1 promoter (<u>#105677</u>, Addgene) and the cytosolic reporter tandem tomato (tdTomato) under the CAG promoter (<u>#83029</u>, Addgene). The plasmids were combined in the following amounts: 7 μL of CAG-Cre (3 μg/μL), 7 μL of stGtACR2 (2.6 μg/μL) and 5 μL of CAG-tdTomato (3.7 μg/μL). 1 μL of 0.03 % fast green dye (Sigma) was added to make a total volume of 20 μL. For each embryo, 1.5 μL of the plasmid-containing mixture was injected intraventricularly. The anode of a 5 mm platinum Tweezertrode (BTX) was then positioned over the dorsal telencephalon external to the uterine muscle and five pulses of 36 V (each pulse 50 ms separated by a 950 ms interval) were delivered using a pulse generator (BTX). After all embryos had been electroporated, the uterine horn was returned to the abdominal cavity, which was filled with warm sterile saline solution. The abdominal wall was closed with Vicryl sutures (Ethicon) and the skin closed with Prolene sutures (Ethicon). Dams were closely monitored during the recovery period. Pups were screened for successful electroporation on the third postnatal day (P3), by illuminating the head with a custom-built light-emitting diode (LED, 525 nm, 30 W) to detect tdTomato fluorescence. The pups were weaned at three weeks of age and maintained until 6-8 weeks for experiments.

### Immunohistochemistry

Mouse brains were fixed via transcardial perfusion of phosphate buffered solution (PBS, 0.1 M, Sigma) and 4% paraformaldehyde solution (PFA, Sigma). Brains were stored for 24 h at 4°C in 4% PFA and then washed and stored in PBS containing 0.05% sodium-azide (Merck). Within a week of perfusion, brains were washed in PBS and mounted onto a microtome (HM650V, Thermo Scientific) before being sectioned into 100 μm thick coronal sections whilst bathed in PBS. Thereafter, sections were washed with fresh PBS before being washed with PBS containing 0.3% triton (Sigma; 0.3% PBST). Sections were then blocked in a solution containing 20% normal goat serum (NGS) in 0.3% PBST for 2 h. The sections were washed with 0.3% PBST after blocking and then incubated at 4°C with the primary antibody solution containing 10% NGS, 0.3% PBST plus the relevant antibody for Cux1 (Santa Cruz, <u>#SC-</u> <u>13024</u>) or PV (Synaptic Systems, <u>#195004</u>). After 24 h, sections were again washed in 0.3% PBST before incubation with the secondary antibody at room temperature (∼25°C) for 2 h in a solution containing 5% NGS, 0.3% PBST and a fluorophore-linked secondary antibody specific for the relevant primary antibody. These included an anti-rabbit 635 for Cux1 labelling (Thermo Fisher, <u>#A-31577</u>) and an anti-guinea pig 488 for PV labelling (Thermo-Fisher, <u>#A-</u> <u>11073</u>). Sections were washed in fresh PBS and mounted on slides with VectaShield (Vectorlabs). Confocal imaging was performed using a LSM 880 micrscope (Zeiss) equipped with 488 nm, 561 nm and 633 nm laser lines. All images were captured using a 20x water-immersion objective (W Plan-Apochromat NA 1.0) with the ZEN software (Zeiss). Image processing was performed in ImageJ software (NIH).

### Acute slice preparation

*In vitro* electrophysiological experiments were performed in acute brain slices prepared from S1 of juvenile/adult mice aged 6-8 weeks, 2-3 weeks after intracerebral viral injection. Mice were first anaesthetized with isoflurane (Zoetis) and decapitated. The brain was then sliced into 350 μm coronal sections using a microtome (HM650V, Thermo Scientific) in carbonated high-sucrose solution maintained at 4°C and consisting of: 185 mM sucrose, 1.2 mM NaH_2_PO4, 25 mM NaHCO_3_, 2.5 mM KCl, 25 mM glucose, 2 mM CaCl_2_ and 2 mM MgCl_2_ (all Merck). The pH was adjusted to 7.3-7.4 (S20, Mettler Toledo) and osmolarity to 300-310 mOsm (5520 Vapro, Wescor). Brain slices were recovered in an incubation chamber maintained at 37°C and containing artificial cerebrospinal fluid (aCSF) consisting of: 120 mM NaCl, 3 mM KCl, 1.2 mM NaH_2_PO_4_, 23 mM NaHCO_3_, 11 mM D-Glucose, 2 mM CaCl_2_ and 2 mM MgCl_2_ (pH 7.3-7.4, mOsm 300-310 mOsM). Slices were recovered for at least 60 min before starting electrophysiological recordings. When required, slices were transferred to a recording chamber and continuously superfused with aCSF bubbled with carbogen gas (32 °C and perfusion speed of 2 ml/min).

### In vitro electrophysiological recordings

All electrophysiological recordings were performed using an Axopatch 700A amplifier with data acquired using Clampex software (Molecular Devices). A HumBug noise eliminator (Digitimer) was used to remove 50 Hz noise. For *in vitro* whole-cell recordings, micropipettes (2-5 MΩ) with a short shaft were prepared from borosilicate glass capillaries (Harvard Aparatus) using a horizontal puller (Sutter) and filled with a K^+^ gluconate based internal solution, which contained (in mM): 126 K-Glu, 4 KCl, 4 Na_2_ATP, 0.3 NaGTP, 10 Na_2_-PhosphoCreatine, 10 HEPES, 0.05 EGTA. For experiments in which intracellular Cl^-^ was varied, this was achieved by adjusting the amount of K-Glu and KCl. For the 20 mM Cl^-^ internal solution, 110 mM of K-Glu and 20 mM KCl were used. For the 70 mM Cl^-^ internal solution, 60 mM K-Glu and 70 KCl were used. All other reagents were maintained the same. For all internal solutions, pH was adjusted to 7.3 using K^+^ hydroxide (Mettler Toledo) and osmolarity was adjusted to 287-293 mOsm (Wescor). Pipettes were back-filled with the internal solution, mounted on a micro-manipulator (Sutter), positive pressure was applied, and the pipette tip was lowered under visual guidance onto the surface of a neuron’s soma within the brain slice. Neurons were visualized using dot contrast and a 60x objective (LUMPlanFI/IR, NA 0.90, Olympus).

To perform perforated patch-clamp recordings, the internal pipette solution was prepared immediately prior to recording by combining a high Cl^-^ (150 mM) solution (in mM: 141 KCl, 9 NaCl, 10 HEPES) heated to 37°C with a stock solution of gramicidin A (4 mg/ml - dissolved in dimethyl sulfoxide, DMSO, Merck) to achieve a final concentration of 80 μg/mL gramicidin^79^. The solution was then vortexed (40 s) and sonicated (20 s), before the pipette was back-filled with the gramicidin solution. Once the gigaseal had formed, perforation was monitored by observing changes in series resistance. Recording protocols were started once the series resistance had stabilized at <50 MΩ. Rupture or breakthrough of the perforation in to whole-cell configuration was detected by sudden and persistent depolarization of the measured E_GABAAR_, consistent with dialysis of the neuron with the high Cl^-^ pipette solution.

stGtACR2 was activated by deliver blue pulses of light via a 1000 μm optic fiber connected to a 473 nm laser (MBL-FN-473-150mW, CNI Laser). The tip of the fiber was positioned at an image plane in the microscope, in the center of the optical axis, and directed into the objective lens via a dichroic mirror. In a subset of experiments, focal GABA (100 μM,Torcis Biosciences) was applied to the soma of the recorded neuron via a micropipette attached to a picospritzer (5 psi, 10 ms, General Valve). To isolate the GABA_A_R response, GABA_B_Rs were blocked by adding the selective antagonist, CGP 55845^80^, at a final concentration of 10 μM in the circulating aCSF. In a subset of experiments, KCC2 was blocked by adding the selective antagonist, VU0463271^81^, at a final concentration of 10 μM in the circulating aCSF.

### In vivo two-photon guided patch-clamp recordings

*In vivo* targeted patch-clamp recordings were performed based on previously published protocols^82–84^. Mice aged 6-8 weeks that had undergone IUE were anaesthetized with an intraperitoneal injection of 25% urethane (1 g/kg in sterile PBS). To counteract adverse events caused by urethane, a bolus of the anticholinergic agent, Glycopyrronium Bromide (0.01 mg/kg) was administered subcutaneously. Local anaesthetic (Marcain 2 mg/kg) was applied intradermally in the scalp and topically in the ears prior to mounting the mouse into the head holding apparatus (Narishige) under a surgical stereoscope (Olympus). The mouse’s body temperature was maintained at 37°C using a heating mat and rectal probe. The animal’s head was shaved and eye-protecting ointment (Viscotears) was applied to both eyes. An incision in the scalp was made using surgical scissors and the area expanded with blunt dissection to expose the skull. The site for the craniotomy over S1 was identified based on the area of maximum tdTomato expression, detected using a custom-built LED (525 nm, 30 W). Tissue adhesive (Vetbond) was applied to fix the surrounding scalp to the skull and to secure cranial sutures. Multiple layers of dental cement (Simplex Rapid) were applied to create a chamber on top of the skull. A 0.5 mm craniotomy was drilled over the marked region using a dental drill (Foredom). The craniotomy was submerged in cortex buffer (containing, in mM: 125 NaCl, 5 KCl, 10 HEPES, 2 MgSO_4_·7H_2_O, 2 CaCl_2_·2H_2_O, 10 Glucose). The bone flap and dura were removed and the animal was then relocated onto the recording apparatus. Neurons were visualized using a custom-built two-photon microscope, which consisted of a modified confocal scan unit (Olympus FV300) coupled to a Ti:Sapphire laser (Newport Spectra-Physics Mai Tai HP; 930 nm, 30 mW) and images were acquired using FluoView software (Olympus).

*In vivo* patch pipettes were prepared from borosilicate glass capillaries (Harvard Aparatus) using a vertical puller (Narishige) to obtain a long shaft and tip size of 5-8 MΩ ^82^. Patch pipettes were filled with the same internal solutions described above, with the addition of 15 mM Alexa Fluor 594 dye (Thermo Fisher) and mounted into an Optopatcher pipette holder (A-M Sytems) on a micro-manipulator (Sutter). stGtACR2 was activated via a 50 μm optic fiber (Thorlabs) embedded within the Optopatcher holder and connected to a 473 nm laser (MBL-FN-473-150mW, CNI Laser). A silver-Cl^-^ ground electrode (Multichannel Systems) was lowered onto the surface of the nearby skull. The recording site was submerged in cortex buffer. Pipettes with positive pressure (200-300 mBar) were lowered onto the brain surface under 4x magnification (Plan N, NA 0.10, Olympus). The pipette was then visualized under 40x magnification (LUMPlanFI/IR NA 0.80, Olympus), inserted into the brain to a depth of 200-300 μm, and the positive pressure was first lowered to lowered to 150-170 mBar and then to 20-30 mBar when the pipette was close to the target neuron. The Alexa dye enabled the pipette to be located under two-photon microscopy (820 nm, 50 mW) and the positive pressure released the Alexa dye into the extracellular space, revealing ‘shadows’ of neuronal cell bodies^85^. A neuron was approached and when consistent changes in electrode resistance appeared and the dimple of the fluorescent dye on the neurons membrane was visible, the positive pressure was released and the holding potential clamped at -70 mV in voltage-clamp mode to assist gigaseal formation. Other conventions were the same as for *in vitro* electrophysiological recordings.

### Determining chloride equilibrium potentials

Voltage-clamp recordings were used to measure the equilibrium potential of stGtACR2 (E_stGtACR2_) and the GABA_A_R (E_GABAAR_). Step protocols involved delivering a series of voltage steps to the neuron, during which a transmembrane Cl^-^ conductance was evoked by either light activation of stGtACR2 or agonist activation of GABA_A_Rs. From a holding potential of -70 mV, the voltage was stepped in 10 mV increments from -130 mV to -30 mV, each step lasted 500 ms, with 10 s between steps, and Cl^-^ conductances were evoked 100 ms following the start of the steps. For analysis purposes, the membrane current was measured immediately before the Cl^-^ conductance (i.e. ‘baseline’ current) and then during the evoked Cl^-^ conductance. The two currents were then superimposed on current-voltage (IV) plots. The voltage was corrected for series resistance effects^86^, by calculating the voltage drop associated with the series resistance and subtracting this from the command voltage. The point at which the two IV curves intersect reflected the equilibrium potential of the evoked Cl^-^ conductance. In addition, the baseline current was used to infer the resting membrane potential (RMP), defined as the membrane potential at which the fitted line was equal to zero.

Voltage ramp protocols were used to monitor dynamic changes in E_stGtACR2_. These ramp protocols afforded frequent measurements of E_stGtACR2_, which could be made before and immediately following a Cl^-^ load. From a holding potential of -70 mV, neurons were subjected to voltage ramps under baseline conditions (‘baseline ramp’) or during light activation of stGtACR2. Each voltage ramp lasted 150 ms and extended from 60 mV below the holding potential to 40 mV above the holding potential (i.e. from -130 mV to -30 mV, at a rate of 0.7 mV/ms). For analysis purposes, the first and last 15 ms of each ramp were excluded to avoid transient currents associated with the pipette capacitance. To isolate the stGtACR2 current associated with each voltage ramp, the baseline ramp current (i.e. without light) was subtracted from the stGtACR2-associated ramp current (i.e. with light). The isolated stGtACR2 current was then plotted on an IV curve, where the voltage was corrected for series resistance effects, as above. E_stGtACR2_ was defined as the voltage at which the isolated stGtACR2 current was equal to zero. Neurons were subjected to a Cl^-^ load by combining a prolonged activation of stGtACR2 (1 s light pulse) and increasing the Cl^-^ driving force by clamping the membrane potential at -20 mV.

### Single-cell RNA sequencing meta-analysis

The levels of KCC2 (SLC12A5) RNA expression in pyramidal neurons and PV interneurons was acquired from published single-cell RNA sequencing datasets from mouse sensory cortex (<u>GSE60361</u>, <u>GSE71585</u>, <u>GSE60361</u>). Each dataset was first reorganized to capture the following variables: single-cell ID, cell type for each single-cell ID, gene tested, and a matrix of the RNA expression level measurements corresponding to each single-cell and gene tested. To normalize the data, the raw KCC2 RNA count for each neuron was divided by the total expression level of all measured genes in that neuron, and the resulting number was multiplied by the mean gene expression level across all cells within the same dataset. The resulting data was then analyzed using the ARTool to perform post-hoc aligned rank transform to measure pairwise comparisons^87, 88^.

### Biostatistics

The majority of the statistical analyses was performed using the Python Scipy library. The only exception was the analysis of time constants which was performed using the one phase decay analysis tool in the GraphPad Prism software package (v8.0.0, Dotmatics). In all analyses, an alpha value (p-value, *p*) of less than 0.05 was considered significant. All data is reported as mean ± standard error of the mean. For comparison statistics, appropriate parametric and non-parametric tests were used.

### Data and code availability

Patch-clamp data was analyzed using Python 3.10 using the pyABF package^89^. All material required to reanalyze data is available from the corresponding authors upon reasonable request.

## Acknowledgements

We would like to thank members of the Akerman Research Group for advice and comments. We thank Joseph Raimondo (University of Cape Town) for his comments on the manuscript. The research leading to these results has received funding from the European Research Council under grant agreement 617670; plus BBSRC project BB/S007938/1 and MRC project MR/S01134X/1. This work was supported by a Shaun Johnson Memorial Scholarship sponsored by the Leverhulme Trust and Mandela Rhodes Foundation (R.J.B.). Wellcome Trust Doctoral Fellowships supported T.D. [222380/Z/21/Z], G.G. [219983/Z/19/Z] and A.C. [102364/Z/13/Z]. J.F.A.P acknowledges support from the European Research Council (ERC-2015-CoG-682422) and Deutsche Forschungsgemeinschaft (DFG, FOR 2143). We thank Lex Kravitz, Luigi Petrucco and Ethan Tyler for sharing their mouse illustrations on the SciDraw opensource platform. Some parts of the figures were created using Biorender under a license granted to R.J.B.

## Author contributions

Conceptualization: R.J.B., C.J.A.

Investigation: R.J.B., T.D., A.C., G.G., J.S.J.

Writing: R.J.B., C.J.A

Supervision: J.F.A.P, A.S., C.J.A.

Funding acquisition: R.J.B., J.F.A.P, A.S., C.J.A.

